# Gene Therapy Mediated Partial Reprogramming Extends Lifespan and Reverses Age-Related Changes in Aged Mice

**DOI:** 10.1101/2023.01.04.522507

**Authors:** Carolina Cano Macip, Rokib Hasan, Victoria Hoznek, Jihyun Kim, Louis E. Metzger, Saumil Sethna, Noah Davidsohn

## Abstract

Aging is a complex process best characterized as the chronic dysregulation of cellular processes leading to deteriorated tissue and organ function. While aging cannot currently be prevented, its impact on lifespan and healthspan in the elderly can potentially be minimized by interventions that aim to return these cellular processes to optimal function. Recent studies have demonstrated that partial reprogramming using the Yamanaka factors (or a subset; *OCT4, SOX2*, and *KLF4; OSK)* can reverse age-related changes *in vitro* and *in vivo*. However, it is still unknown whether the Yamanaka factors (or a subset) are capable of extending the lifespan of aged wild type mice. Here, we show that systemically delivered AAVs, encoding an inducible OSK system, in 124-week-old mice extends the median remaining lifespan by 109% over wild-type controls and enhances several health parameters. Importantly, we observed a significant improvement in frailty scores indicating that we were able to improve the healthspan along with increasing the lifespan. Furthermore, in human keratinocytes expressing exogenous OSK, we observed significant epigenetic markers of age-reversal, suggesting a potential reregulation of genetic networks to a younger, potentially healthier state. Together, these results may have important implications for the development of partial reprogramming interventions to reverse age-associated diseases in the elderly.

## Introduction

The world’s population is growing older, with a doubling of the median age from 1900 to 2020, leading to increased societal burden (Partridge et al., 2018). Aging is the strongest risk factor for most common human diseases (Partridge, 2014), hence it is imperative to identify anti-aging interventions to delay or even potentially reverse the aging process. Increasing longevity has historically referred to extending the ‘lifespan’ of an organism through various interventions such as public health policies (Ray M Merrill, 2014), caloric restriction (de Cabo et al., 2014; López-Otín et al., 2013; Swindell, 2012), or through pharmaceutical interventions (Blagosklonny, 2019; Glossmann and Lutz, 2019). One potential pitfall of increasing longevity is that it may not necessarily improve quality of life or healthspan. For example, an organism may live longer but still experience age-related diseases and physiological decline, albeit on a longer timescale. Age reversal, on the other hand, refers to the process of restoring an organism to a younger state, abrogating the effects of aging at the cellular level, and consequently increasing both healthspan and lifespan.

The other pitfall of longevity research is cycle-time. For assessment and development of potentially efficacious interventions, it would necessitate waiting for the organism to die. Many groups are working to elucidate biomarkers that are sensitive and correlate reliably with increased lifespan (Horvath, 2013; Horvath and Raj, 2018; Hsu et al., 2020), yet the current gold standard remains ‘time to death’. This readout works well for short-lived multicellular model organisms such as *C. elegans* (∼3 weeks) (Zhang et al., 2020) and *D. melanogaster* (∼70 days) (Piper and Partridge, 2018). At the mouse level, testing anti-aging interventions can take 0.5 to 3 years.

Using a cocktail of transcription factors, *OCT4* (O), *SOX2* (S), *KLF4* (K), and *c-MYC* (M), collectively known as OSKM or Yamanaka factors, seminal studies showed that somatic cells can be reversed to a pluripotent state (Takahashi and Yamanaka, 2006), thereby reversing a long-held paradigm of unidirectional differentiation. By short or cyclic induction of the Yamanaka factors in transgenic mice, investigators have demonstrated age extension in progeroid mice. These transgenic mouse models encoded a polycistronic OSKM cassette driven by a reverse tetracycline transactivator (rtTA) (4F mice); cyclic administration of doxycycline led to partial reprogramming without teratoma formation. This paradigm partially ameliorated aging phenotypes and extended the lifespan in the 4F-progeroid model. The study further showed that the epigenetic profile assessed by epigenetic methylation clocks (Horvath, 2013; Horvath and Raj, 2018) of tissues, correlated with improved function. (Ocampo et al., 2016). Another study demonstrated that short induction of OSKM in a myocardial infarction model alleviated myocardial damage and improved cardiac function (Chen et al., 2021).

The translation of these proof-of-concept genetic studies toward therapeutic interventions is to benefit the increasingly large aging population. In support of this endeavor, we have generated a systemically delivered two-part AAV9 system with doxycycline-inducible OSK. By cyclic induction of this system, we observed a 109% increase in median remaining life in 124-week-old OSK treated mice relative to doxycycline treated control mice. Moreover, we showed that constitutive OSK expression in human keratinocytes leads to profound age-reversal as assessed by methylation clocks.

## Results

Transgenic mouse models are not suitable to enable translation of therapeutic strategies to humans for age reversal, hence we used an AAV system to systemically deliver OSK. Secondly, young humans are not the target population for age-reversal therapeutics, hence we chose extremely old mice (124 weeks) as a model system for improved translatability. Wild type C57BL6/J mice have a median lifespan of ∼129 weeks (Yuan et al., 2012), equivalent to ∼80 years in humans (Ackert-Bicknell et al., 2015). We drove inducible OSK expression in 124-week mice (∼77 years in human age) using a two part AAV system, where one vector carried a constitutively expressed rtTa and the other vector contained a polycistronic OSK expression cassette driven by doxycycline responsive TRE promoter (Lu et al., 2020) (**Fig. 1a**). We selected AAV9 capsid to ensure maximal distribution to most tissues (Inagaki et al., 2006). We injected 124-week-old WT C57BL6/J mice retro-orbitally (RO) with 100 μl containing either PBS (formulation buffer) or 1E12 vg of each vector for a total dose of ∼6E13 vg/kg. We initiated the doxycycline induction for both the control and AAV administered groups the day after injections and alternated weekly on/off cycles for the remainder of the animals’ lives (details in **Methods)**. Intriguingly, we observed a 109% extension in median remaining life in response to OSK expression (control mice had 8.86 weeks of life remaining vs. 18.5 weeks for TRE-*OSK* mice). Doxycycline-treated control mice had a median lifespan of ∼133 weeks, while the TRE-*OSK* mice had a median lifespan of 142.5 weeks (**Fig. 1b, c and Supplementary Fig. 1**). We further compared the control doxycycline treated mice to the historical published data for Bl6/J mice (Yuan et al., 2012) and available via the mouse phenome database (https://phenome.jax.org/projects/Yuan2); we found no significant differences in median survival, suggesting that doxycycline alone had no adverse nor advantageous effects (**Fig. 1b**).

**Fig. 1:**
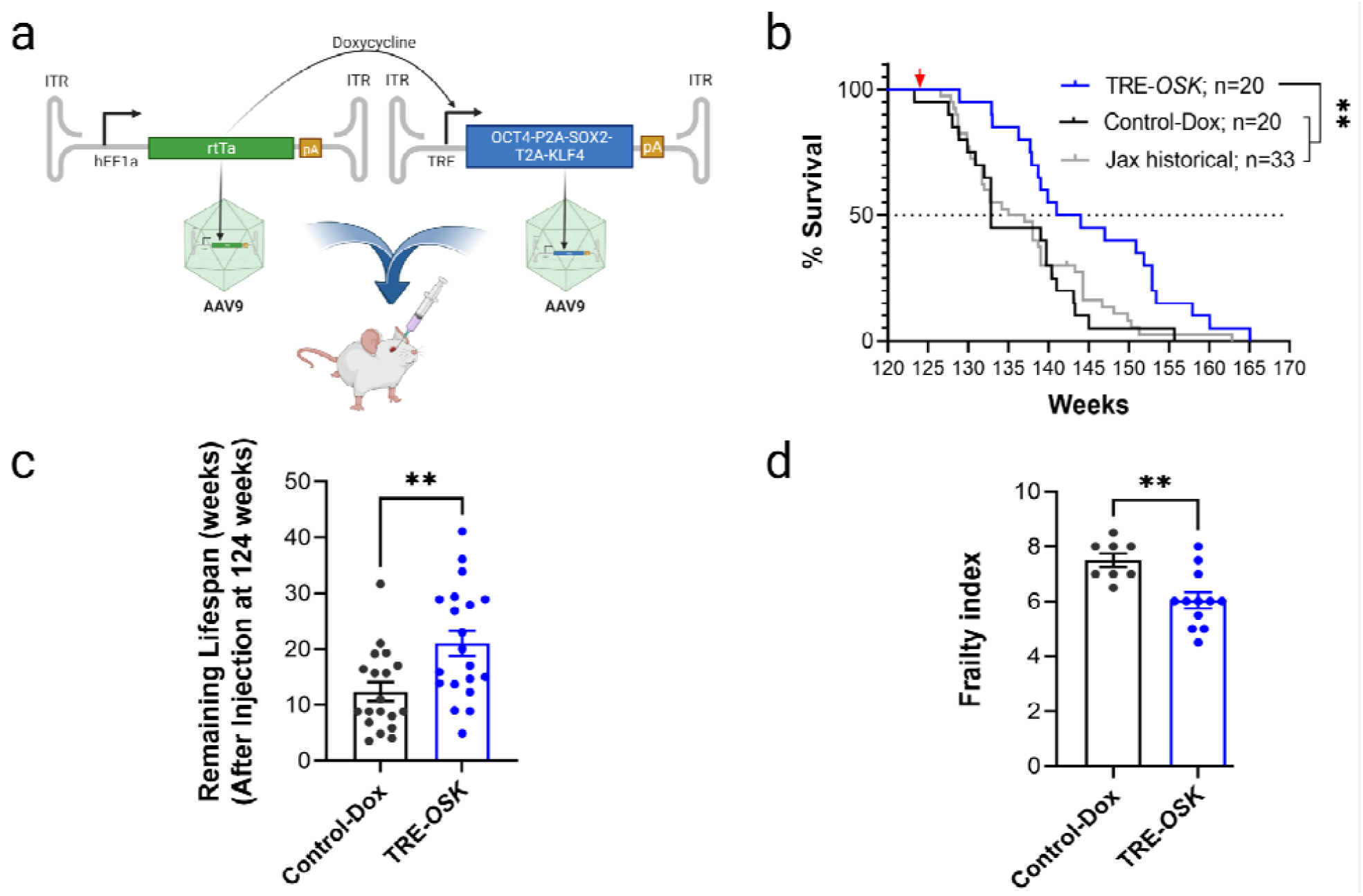
Partial reprogramming with TRE-*OSK* leads to increased lifespan and improved frailty scores in very old mice. **a**. Schematic of the constructs, virus, and injection route used in the study. **b** Kaplan Meier curves for 124-week WT mice injected with AAV9.TRE-*OSK* and AAV9.hEF1□-*rtTA4* (both 1E12 vg/animal) via the retro-orbital route, and induced with one week on/off doxycycline paradigm (TRE-*OSK*) showed median lifespan extension of remaining life by 109% compared to either doxycycline treated control animals (Control-Dox) or to historical published Jax data for Bl6/J mice (Jax historical). Red arrow indicates AAV injections. Mantel Cox Log rank test, ** *p*<0.05. **c**. Graph shows remaining lifespan of individual mice (after injections at week 124) for data shown in **b**. Two-tailed unpaired *t*-test; ** *p*<0.05. **d**. Frailty index (FI), the compound score of 28 different health parameters (range 0-1 in 0.5 increments), showed significant reduction in FI for TRE-*OSK* mice at 142 weeks of age (18 weeks after injections) as compared to Control-Dox mice. Student’s unpaired *t*-test, ** *p*<0.05.

Aging is associated with an increased susceptibility to adverse health outcomes which can be captured by clinicians using a frailty index (FI), where people are scored based on a subset of age-related health deficits. High compound scores reflect a frail state and increased susceptibility to poor health outcomes (Searle et al., 2008). A similar index can be used in mice to assess aging and effects of aging interventions (Heinze-Milne et al., 2019). We observed a significant reduction in the FI from 7.5 points for doxycycline treated control mice to 6 points for TRE-*OSK* mice (**Fig. 1d**), suggesting that increased lifespan correlated to overall better health of the animals.

Molecular measures of cellular and tissue health have been developed based on methylation patterns of genomic DNA. “Epigenetic age”, a well characterized and established aging biomarker, can be calculated using these methylation patterns. Such epigenetic clock biomarkers decouple chronological age (Bell et al., 2019) from the functional state of the cells or tissues, while correlating better to aging, disease state(s), and health outcomes (Bell et al., 2019; Durso et al., 2017; Fransquet et al., 2019; Xiao et al., 2021). We isolated DNA from heart and liver tissue from control and TRE-*OSK* treated mice at time of death and found that the Lifespan Uber Clock (LUC) (Browder et al., 2022; Haghani et al., 2022) for both liver and heart trended towards reduced epigenetic age (**Fig. 2a**).

**Fig. 2:**
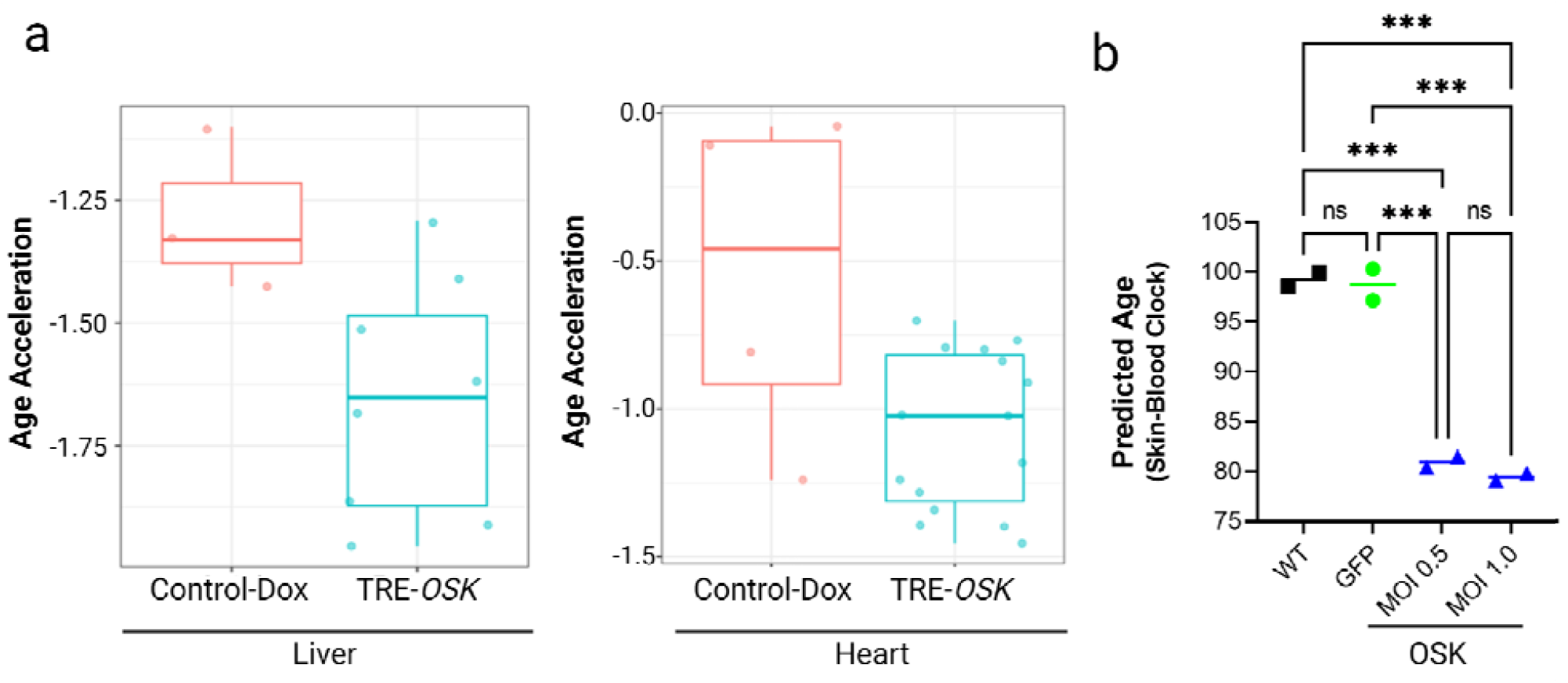
Partial reprogramming with TRE-*OSK* in human keratinocytes leads to age-reversal as assessed by genomic methylation state. **a**. Measurement of DNA methylation age acceleration in liver (*left* panel) and heart (*right* panel) from control doxycycline treated mice (Control-Dox) or TRE-*OSK* mice using LUC epigenetic clocks trained on the indicated tissues. **b**. Human keratinocytes transduced with lentivirus at 2 different MOIs (0.5 and 1.0) expressing OSK showed epigenetic age reversal as compared to control GFP transduced or non-transduced (WT) cells. n=2 technical repeats for each group. One-Way ANOVA with Holm-Šídák’s multiple comparisons test. ns - not significant; *** *p*<0.001.

To assess the effects of OSK over-expression in a cellular system, we expressed OSK in HEK001 keratinocytes isolated from the scalp of a 65-year-old male patient. We confirmed, by immunoblot, the exogenous expression of OSK in these keratinocytes transduced with lentivirus (**Supplementary Fig. 2**). Next, we found significant epigenetic age reversal in keratinocytes treated with OSK as compared to either untransduced or GFP transduced cells (**Fig. 2b**). Taken together, our mouse and keratinocytes data suggest that AAV-mediated gene therapy delivering OSK increases lifespan in mice with improved health parameters and reverses biomarkers of aging in human cells

## Discussion

In modern societies, aging is the highest risk factor associated with most diseases and mortality (Partridge, 2014). The goals of regenerative medicine are to improve tissue and organ function and to correct disease states. Cellular rejuvenation via partial reprogramming has been shown to be a promising avenue to achieve the goals of regenerative medicine. Here we show that in human cells, exogenous expression of a OSK leads to profound age reversal as observed by the restoration of genomic methylation patterns to those characteristic of younger cells, a validated hallmark of chronological age-reversal (Bell et al., 2019; Haghani et al., 2022; Xiao et al., 2021). To our knowledge we have shown for the first time an extension of remaining median lifespan in extremely old WT C57BL6/J mice concomitantly with improved health outcomes as a consequence of a systemic AAV-based therapy. Experiments to assess the epigenetic programming hallmarks in specific tissues, along with thorough analysis of the RNA profiles from these studies will be required to make broader conclusions as to which pathways are reprogrammed to a more youthful state.

We assessed whether *in vivo* partial cellular rejuvenation is sufficient to extend lifespan and healthspan in a relevantly old population and to remove a major barrier to the systemic delivery of three Yamanaka factors within a single vector. Investigators have hitherto shown transduction of specific organs with combinations of OSK or OSKM, but with each encapsulated in a separate vector (Senís et al., 2018). For therapeutic development in humans, having three separate vectors significantly increases the complexity of manufacturing, drug product specifications, and administration protocols for clinical development. Based on our novel proof-of-concept studies in an extremely aged mouse population (equivalent to >80 years of age in humans) and previous studies in younger mice (Browder et al., 2022; Lu et al., 2020; Ocampo et al., 2016), we envision therapeutic rejuvenation in aging humans, first in specific age-related disease settings and later for therapeutic healthspan and lifespan extension. Teratoma formation has been observed in partially reprogrammed animals, particularly when c-Myc is used in the partial rejuvenation cocktail (Abad et al., 2013; Ocampo et al., 2016; Ohnishi et al., 2014; Senís et al., 2018). Although poorly invasive and poorly metastatic, teratoma formation is unlikely to be accepted by the FDA, hence tight control of the partial rejuvenation factors will be a key attribute for safe and efficacious rejuvenation therapies. We did not observe any gross teratoma formation in our cyclically induced OSK paradigm. These observations, along with recent advances in vector development and optimization, tissue specific promoters, and inducible systems (Domenger and Grimm, 2019; Li and Samulski, 2020), engender cautious optimism that a partial rejuvenation therapy can be safely delivered in humans. Prudent and thorough monitoring studies in large animals will be required to assess the safety and efficacy of partial rejuvenation studies.

## Materials and Methods

### Vector and AAV generation

Constructs containing tetracycline-responsive element version 3 (TRE3) promoter driving the expression of human *OCT3/4, SOX2*, and *KLF4* from a polycistronic transcript (TRE3-*OSK*) and second construct encoding rtTA version 4 driven by hEf1a promoter (hEf1a-*rtTA4*) were generated by Genscript (Piscataway, NJ) as reported previously (Lu et al., 2020). The constructs were packaged in AAV9 capsid to generate AAV9.TRE3-*OSK*-SV40pA (1.556 E13 vg/ml) and AAV9-hEf1a-*rtTA4*-Sv40pA (1.88 E12 vg/ml) by SignaGen (Fredrick, MD).

### Mouse studies and frailty scores

Mouse experiments were performed at Jax labs (AUS protocol #19063). Male C57BL6/J (JAX Stock# 000664) mice aged to 124 weeks were injected with the two viruses described above; each 1E12 vg/mouse (in 100 μl volume) via retro-orbital route. Control mice were injected with 100 μl formulation buffer (PBS). Doxycycline induction was performed one week on/one week off for the duration of the study, by providing 2 mg/ml final concentration of doxycycline in drinking water. Control mice received doxycycline in water at the same concentration as the vector injected mice. The euthanasia criteria was as follows; a rapid or sustained deterioration in health status resulting in a body condition score (BCS) of ≤ 2; tumors or other masses that become ulcerated or interfere with the ability of the animal to eat, drink, or ambulate; any prolapsed organs that cannot be reduced and/or become ulcerated and/or necrotic; any other condition that interferes with ability to reach or consume adequate amounts of food or water. Mice were individually weighed and assessed across 28 different variables including physical, physiological, and innate reflex conditions, including simple sensorial and motor tests, body temperature and overall body condition assessment. A Frailty Index score (FI) (Heinze-Milne et al., 2019) is calculated per mouse by adding all individual scores (excluding body temperature and weight) together detailed in **Supplementary Table 1**.

### DNA extraction from tissues and methylation studies

Mice that were healthy and euthanized at the end of the study were selected for methylation studies. Tissue from the liver and the heart was extracted using the DNeasy Blood and Tissue Kit (Quiagen), following the manufacturer’s protocol.

| RO injections | Age (weeks) | Liver | Heart |
| --- | --- | --- | --- |
| TRE-OSK+rtTA | 153.4 | yes | yes |
| TRE-OSK+rtTA | 151.9 | yes | yes |
| TRE-OSK+rtTA | 141.0 | yes | yes |
| TRE-OSK+rtTA | 152.9 | yes | yes |
| TRE-OSK+rtTA | 152.9 | yes | yes |
| TRE-OSK+rtTA | 164.7 | yes | yes |
| TRE-OSK+rtTA | 141.0 | yes | yes |
| TRE-OSK+rtTA | 139.9 | yes | yes |
| Control | 141.0 | yes | yes |
| Control | 143.3 | yes | yes |
| Control | 139.0 | yes | yes |

Methylation analysis on above extracted DNA was performed by the Clock Foundation (Torrence, CA). LUC clock algorithm and analysis has been described previously (Browder et al., 2022; Haghani et al., 2022).

### Lentivirus Production and Keratinocyte Transduction

Plasmids encoding polycistronic OSK driven by human EF1□ promoter, PsPax2 and PmD2.G were co-transfected into HEK293T cells with PEI and Opti-MEM (Gibco). The next day, the media was replaced with harvest media containing DMEM, 15% FBS, and 1% PenStrep (Gibco). Supernatant was collected on days 3 and 4, filtered through a 0.45 polyethersulfone membrane, 1x volume of Lenti-X concentrator (Takara Bio) was combined with 3x volumes of clarified viral media and stored at 4°C overnight. Viral media was spun at 1,500 *x g* for 45 minutes, pellet was resuspended in DMEM. Virus was titered using Lenti-X GoStix Plus (Takara Bio). Lentivirus encoding GFP was purchased by VectorBuilder (Chicago, IL).

Lentivirus containing medium was added dropwise to HEK001 (ATCC CRL-2404) passage 110 containing 8ug/ml of polybrene (Millipore-Sigma) at 2 different MOIs: 0.5 and 1.0. Puromycin at 1 ng/μl (Millipore-Sigma) was added on day 2 for selection. Surviving cells were expanded and maintained with puromycin, changing media every 3 days and splitting as necessary. On day 23 after selection, cells were collected for immunoblot and methylation analysis.

### Immunoblot Analysis

Protein from cells described above was extracted on ice using Lysis Buffer (CST) with 1mM PMSF and protease inhibitor cocktail. Cell lysis mixture was spun for 10 minutes at 14,000g/4°C, supernatant was collected and protein was quantified using Pierce Rapid Gold BCA Protein Assay Kit (Thermo Fisher). Equal amounts of protein were loaded in a 4-15% polyacrylamide gel (Thermo Fisher) and transferred to PVDF membranes and blocked with 5% dry-milk in TBST for 1 hour at room temperature. Membranes were incubated overnight at 4°C with primary antibodies. The following day, membranes were incubated in a secondary antibody conjugated to HRP for 1 hour at room temperature and developed with ECL Prime Western Blotting Detection Reagent (Civita Life Sciences).

Antibodies Used for immunoblot

| Name | Cat # | Manufacturer | Dilution | Host species |
| --- | --- | --- | --- | --- |
| Oct4 | Ab181557 | Abcam | 1:1000 | rabbit |
| Sox2 | Ab92494 | Abcam | 1:1000 | rabbit |
| Klf 4 | PA5-20897 | Thermo Fisher | 1:1000 | rabbit |
| GAPDH-AF488 | MAB374-AF488 | Millipore Sigma | 1:1000 | mouse |
| Rabbit IgG-HRP | Ab6721 | Abcam | 1: 2500 | goat |

## Acknowledgements

We would like to thank Bailey Bonet, Mehdi Doroudchi, Chris Liu, Muralidhar Reddivari, and Chris de Solis for critical reading and suggestions of the manuscript. We would also like to thank Bobby Brooke at the Clock Foundation for the time and effort spent explaining the epigenetic clock results and analysis included in this paper. Fig. 1a was generated using BioRender. This material is based upon work supported by The United States Special Operations Command under Contract No. H9240521C0015. Any opinions, findings and conclusions or recommendations expressed in this material are those of the author(s) and do not necessarily reflect the views of The United States Special Operations Command.

## Competing interests

All authors performed the work while employed at Rejuvenate Bio Inc.

**Supplementary Fig. 1:**
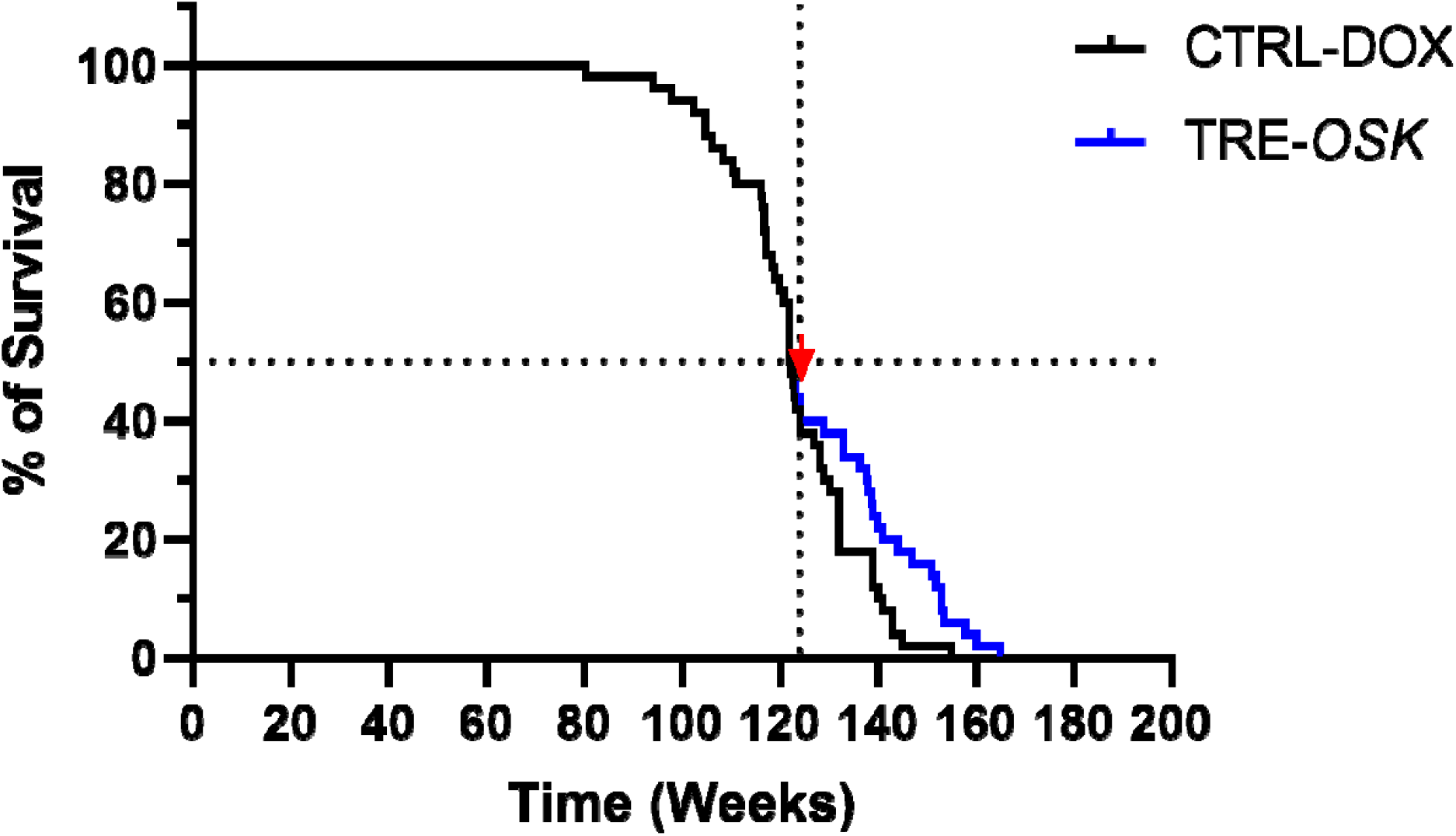
Overall survival proportion curves for control mice and TRE-*OSK* mice over the entire lifespan. Survival proportions for the whole time course for data shown in Fig. 1.

**Supplementary Fig. 2:**
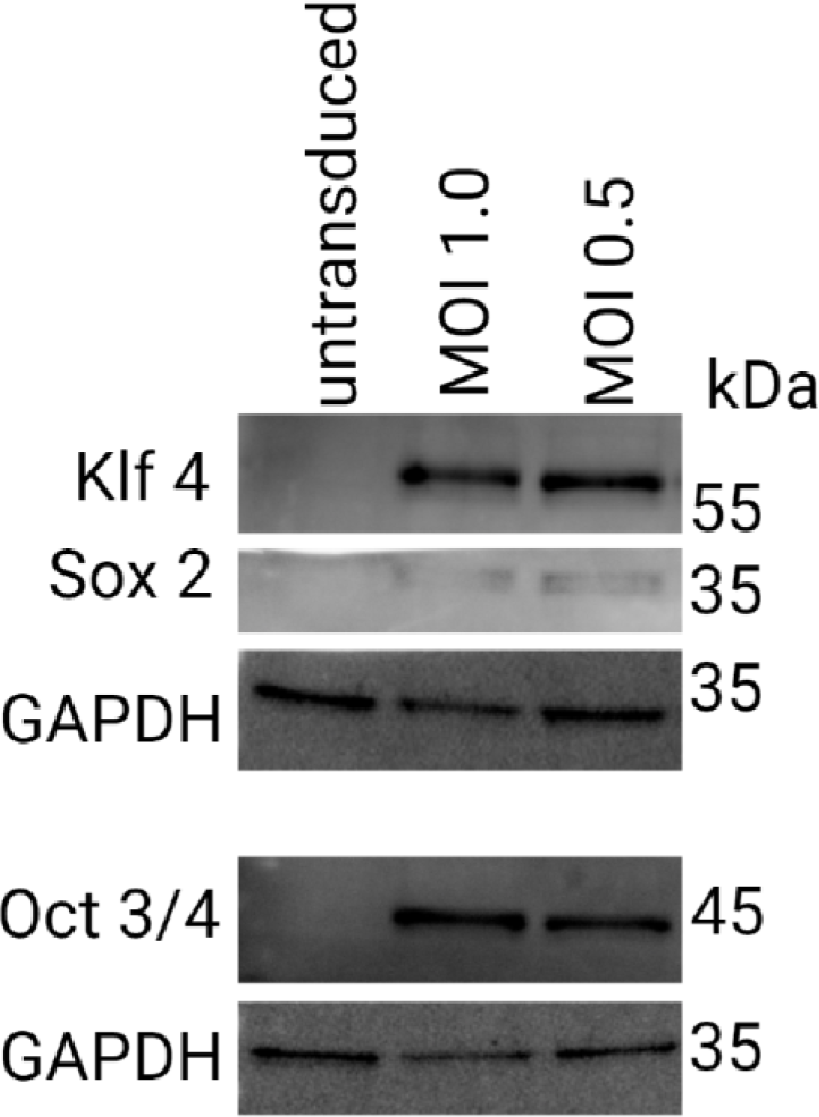
Immunoblots for OSK expression in keratinocytes.

